# Positive selection of ORF3a and ORF8 genes drives the evolution of SARS-CoV-2 during the 2020 COVID-19 pandemic

**DOI:** 10.1101/2020.04.10.035964

**Authors:** Lauro Velazquez-Salinas, Selene Zarate, Samantha Eberl, Douglas P. Gladue, Isabel Novella, Manuel V. Borca

**Affiliations:** Foreign Animal Disease Research Unit, USDA/ARS Plum Island Animal Disease Center, PO Box 848, Greenport NY 11944, USA; College of Veterinary Medicine, Kansas State University, Manhattan, KS 66506, USA; Posgrado en Ciencias Genómicas, Universidad Autónoma de la Ciudad de Mexico, Mexico City, Mexico; Central Connecticut State University, New Britain, CT 06053; Independent researcher

**Keywords:** Novel coronavirus, pandemic, evolution, epistasis, positive selection, COVID-19, SARS-CoV2, phylogenetics

## Abstract

In this study, we analyzed full-length SARS-CoV-2 genomes from multiple countries to determine early trends in the evolutionary dynamics of the novel COVID-19 pandemic. Results indicated SARS-CoV-2 evolved early into at least three phylogenetic groups, characterized by positive selection at specific residues of the accessory proteins OFR3a and ORF8a. We also report evidence of epistatic interactions among sites in the genome that may be important in the generation of variants adapted to humans. These observations might impact not only public health, but also suggest more studies are needed to understand the genetic mechanisms that may affect the development of therapeutic and preventive tools, like antivirals and vaccines.

## Introduction

The first case of pneumonia confirmed to be caused by the novel virus SARS-CoV-2 was in a 41-year-old man in Wuhan, Hubei province, China on 31/December/2019 (Wu et al., 2020). As of 9/April/2020, the World Health Organization (WHO) has confirmed 1,439,516 cases, 85,711 deaths, and the presence of COVID-19 in 209 countries, areas or territories. Of the confirmed cases, 71% are from seven countries: United States of America (395,030), Spain (146, 690), Italy (139, 422), Germany (108,202), China (83,249), France (81,095), and Iran (66,220) (https://www.who.int/emergencies/diseases/novel-coronavirus-2019). As of the writing of this report, the number of COVID-19 cases continue to increase worldwide, with multiple epicenters.

The International Committee on Taxonomy of Viruses (ICTV) initially named this pathogen 2019-nCoV (also referred to as COVID-19 by WHO) and included it within the *Coronaviridae* viral family (Coronaviridae Study Group of the International Committee on Taxonomy of, 2020). Later, based on the close phylogenetic relationship of COVID-19 with other human and bat severe acute respiratory syndrome coronaviruses (SARS-CoVs), ICTV renamed the virus as SARS-CoV-2 (Coronaviridae Study Group of the International Committee on Taxonomy of, 2020).

The *Coronaviridae* family encompasses a group of single-stranded, positive-sense RNA viruses with a genome length varying between 27 and 32 kb. These are zoonotic viruses with the potential to infect humans and animals. Coronaviruses may cause acute and chronic respiratory, enteric, and central nervous system infections(Phan et al., 2018, Weiss and Navas-Martin, 2005). In the case of SARS-CoV-2, a meta-analysis of 50,466 patients indicate that fever and cough are the most common symptoms (95% CI: 81.8-94.5% and 65.7-78.2%, respectively) (Sun et al., 2020). The disease may worsen, and the percentages of severe cases and fatality rate vary between 12.7-24.3% and 2.7-6.1% (95% CI), respectively (Sun et al., 2020).

The genome organization of SARS-CoV-2 is similar to viruses from the genus *Betacoronavirus*, one of the four genera included in the *Coronaviridae* subfamily Orthocoronavirinae. The ∼29,903 nucleotide (nt) genome is organized as follows, 5’ to 3’: replicase ORF1ab, S (encoding the structural spike glycoprotein), ORF3a (ORF3a protein), E (structural envelope protein), M (structural membrane glycoprotein), ORF6 (ORF6 protein), ORF7a (ORF7a protein), ORF7b (ORF7b protein), ORF8 (ORF8 protein), N (structural nucleocapsid phosphoprotein), and ORF10 (ORF10 protein). ORF1ab (∼21,291 nt) encodes sixteen non-structural proteins: leader protein, nsp2, nsp3, nsp4, 3C-like proteinase, nsp6, nsp7, nsp8, nsp9, nsp10, RNA-dependent RNA polymerase, helicase, 3’-to 5’ exonuclease, endoRNAse, 2’-o-ribose methyltransferase, and nsp11(Wu et al., 2020).

Much speculation regarding the origin of SARS-CoV-2 emanates from unfounded theories, such as a man-made laboratory origin; however, a recent study supports the hypothesis that SARS-CoV-2 was the result of cross-species transmission followed by natural selection in the novel human host (Andersen et al., 2020). This hypothesis is strongly supported by studies examining amino acid differences between SARS-CoV-2 and some phylogenetically related betacoronaviruses (e.g., Bat-RatG13 isolate and the human SARS-CoV isolate Urbani) at the receptor-binding domain (RBD) of the spike protein, where such differences seem to increase the ability of SARS-CoV-2 to bind to the human receptor angiotensin-converting enzyme 2 (ACE2) (Andersen et al., 2020). This increased affinity for binding ACE2 might help to explain the infectiousness of SARS-CoV-2 in human populations (Wan et al., 2020).

Considering the extraordinary plasticity shown by other human viral RNA pathogens, for example HIV-1, Influenza viruses, SARS-CoV, and hepatitis C virus, to undergo adaptative changes to evade innate and adaptive immune responses, develop drug resistance or establish an infection in a new host (Frost et al., 2018), multiple questions arise regarding the adaptative changes that SARS-CoV-2 has undergone during the pandemic. SARS-CoV-2 has spread throughout many countries resulting in the infection of people with diverse immunological backgrounds and demographics (age, sex, environmental conditions, etc) that potentially impose significant selective pressures on SARS-CoV-2.

Here, we evaluate the phylogenetic and evolutionary dynamics of SARS-CoV-2 during the early phase of the COVID-19 pandemic. Using different analyses based on a codon-based phylogenetic framework, we identified critical sites in the genome undergoing positive selection, which might favor viral divergence and emergence of multiple viral variants. Our findings are discussed in terms of the potential effects that the rapid evolution of SARS-CoV-2 might have on public health.

## Materials and methods

### Data collection

Eighty-six full-length SARS-CoV-2 genomes representing early viral isolates from patients living in diverse geographic regions were used for this study. Viral sequences, downloaded from the NCBI GenBank database on 06/03/2020, represent the total number of full-length viral genomes at the time that the analysis was conducted (Figure 1).

**Figure 1.**
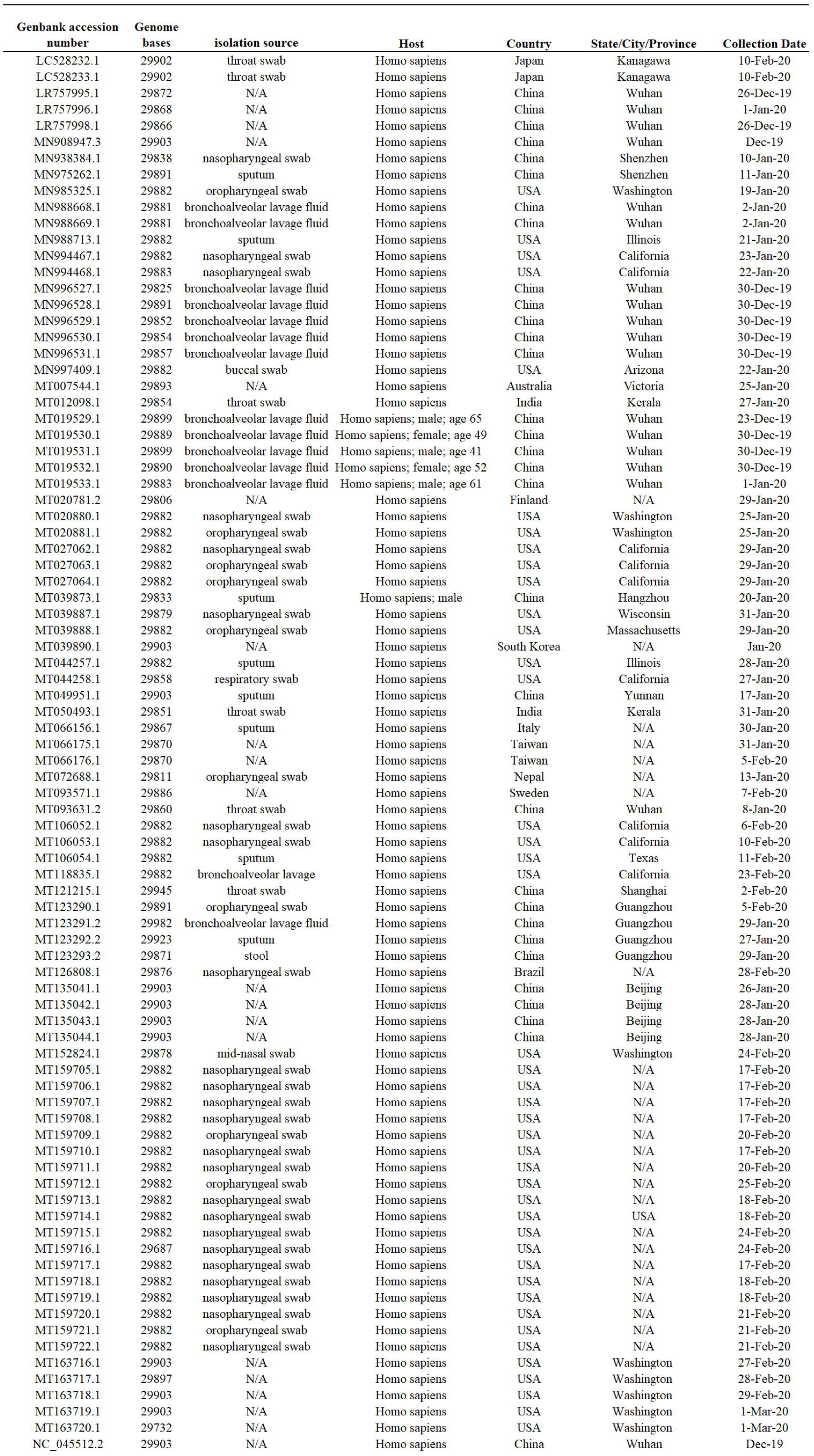
Sample summary. Description of the 86 SARS-Cov-2 full-length genome sequences included in this study. All sequences were obtained form I from the GenBank database, accession number, genome length, isolate name, source, host and country of origin are provided. N/A indicates information not available.

### Phylogenetic dynamic analysis

Two phylogeographic trees were reconstructed using the programs BEAST2.4.3, BEAUti, and TreeAnnotator (Drummond et al., 2012), introducing the country/region as a trait. The Markov Chain Monte Carlo was run for 25 million generations, using the HKY85 substitution model and a gamma distribution with four categories as the site heterogeneity model. The resulting file was analyzed with Tracer 1.6 to check for convergence and to determine the Burnin proportion (Rambaut et al., 2018). Finally, TreeAnnotator was used to build the maximum clade credibility tree, which was visualized with FigTree 1.4.3 (http://tree.bio.ed.ac.uk/software/figtree). Additionally, the evolutionary rate of SARS-CoV-2, expressed as substitutions/sites/year, was calculated using the same methodology but using sampling date as a trait.

### Pairwise distance calculations

Nucleotide and amino acid pairwise distance calculations among SARS-CoV-2 sequences were conducted using the SSE 1.3 Sequence Distances program (Simmonds, 2012). For this purpose, a sliding window of 50 nucleotides (nt), with a shift of 25 nt, was used to determine pairwise distances. Additionally, p-distances in nucleotide and amino acid sequences between phylogenetic groups were calculated using MEGA 7 (Kumar et al., 2016).

### Evolutionary rate per site analysis

Mean (relative) evolutionary rates for each site in the alignment were estimated under the General Time Reversible model, including all 3 codon positions. These rates were scaled considering the average evolutionary rate across all sites is 1. This means that sites showing a rate <1 are evolving slower than average, and those with a rate >1 are evolving faster than average. This analysis was conducted using MEGA 7 (Kumar et al., 2016).

### Inference of selective pressures

Since natural selection can be manifested as different modes (diversifying, directional or purifying), we used a combination of different evolutionary analyses to enhance the detection of relevant sites in the genome of SARS-CoV-2 experiencing diversifying (positive) and purifying (negative) selection: Single Likelihood Ancestor Counting (SLAC) (Kosakovsky Pond and Frost, 2005), Fixed Effects Likelihood (FEL) (Kosakovsky Pond and Frost, 2005), Mixed Effects Model of Evolution (MEME) (Murrell et al., 2012), and Fast Unbiased Bayesian Approximation (FUBAR) (Murrell et al., 2013). These methods use a maximum likelihood or Bayesian approach (FUBAR) to infer nonsynonymous (dN) and synonymous (dS) substitution rates on a per site basis for given coding alignment and corresponding phylogeny (Weaver et al., 2018). SLAC, FEL and FUBAR were methods used to identify sites experiencing pervasive diversifying or purifying selection, while MEME was used to detect sites experiencing episodic diversifying selection.

The presence of recombination in the sequence dataset potentially affecting the detection of positive selection was assessed using the algorithm GARD (Kosakovsky Pond et al., 2006). All methods were performed on the adaptive evolution server Datamonkey 2.0 (Weaver et al., 2018).

Evidence of directional selection was assessed on amino acid sequences using the Directional Evolution of Protein Sequences (DEPS) method, implemented on the Datamonkey webserver (classic) (Delport et al., 2010). This method is a model-based phylogenetic maximum likelihood test that looks for evidence of preferential substitution toward a given residue at individual positions of a protein alignment (Kosakovsky Pond et al., 2008).

### Coevolution analysis

Evidence of coevolution among different sites in the SARS-CoV-2 genome was evaluated using the method Bayesian Graphical Models for co-evolving sites (BGM) (Poon et al., 2007). This method detects coevolutionary interactions between amino acids in a protein, where amino acid substitutions are mapped to branches in the phylogenetic tree.

### Blosum 62 substitution matrix (BSM62)

BSM62 was used to infer the biological significance of amino acid replacements found during the evolutionary analysis of SARS-CoV-2, where positive values reflect that the substitution is most likely a product of random substitution, while negative values are indicative of selection (Henikoff and Henikoff, 1992).

## Results

### Phylogenetic dynamics of SARS-CoV-2

To evaluate potential divergence events of SARS-CoV-2, indicating the rise of new variants early during the pandemic, we reconstructed the evolution of SARS-CoV-2 using full-length genome sequences of viruses collected between late December of 2019 and early March of 2020 from patients infected in different countries around the world. The results of the phylogenetic analysis demonstrate the rapid divergence of SARS-CoV-2 into three distinct phylogenetic groups, differentiated only by a few changes at the nucleotide and amino acid levels (Figure 2A and 2B).

**Figure 2.**
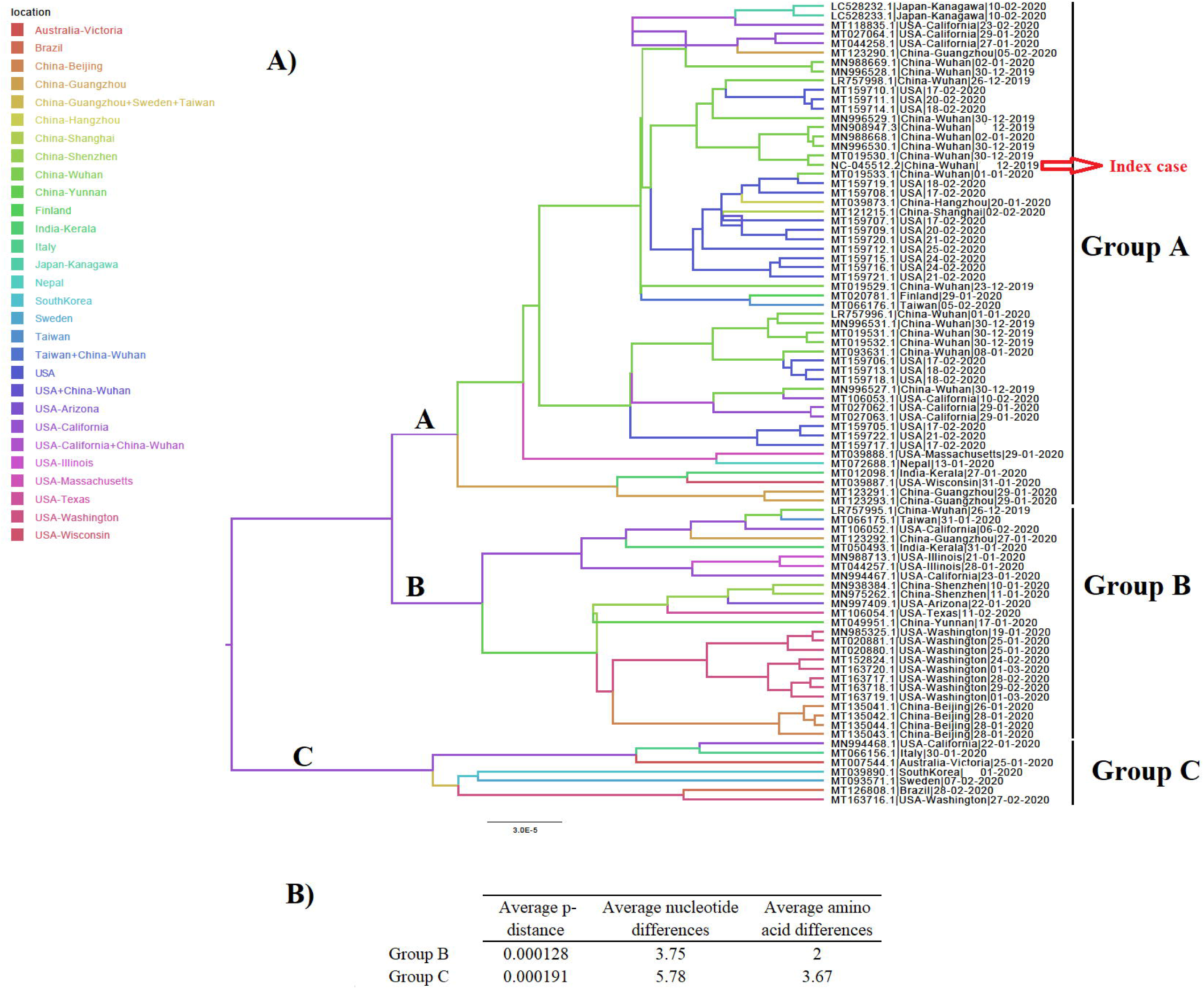
Phylogeny of SARS-Cov-2. A) A MCC tree reconstructed using 86 SARS-Cov-2 full-length genomes collected from patients naturally infected at different countries, showing the existence of 3 phylogenetic groups (A, B, C). Branches in the tree are colored based on the geographic location of each isolate. Presumed index case is pointed out with a red arrow. B) Average p-distance, and average nucleotide and amino acid differences between group A and groups B and C sequences, respectively. Analysis was conducted using the software MEGA 7.

Group A includes the deduced ancestral sequence (NC_045512.2) obtained from the index case in Wuhan, China, and reported to the WHO on 31/December/ 2019, as well as multiple viral isolates from different Chinese provinces. The position of these sequences among multiple branches within the Group A cluster suggests the emergence of multiple viral variants in China, especially from Wuhan before the start of the global pandemic. Furthermore, the basal branch position of some of these variants indicates that they were the ancestors of viral isolates obtained from patients in the USA, Japan, Finland, Taiwan, Nepal, and India between January and February 2020.

Similarly, in the Group B cluster, we found viral isolates from multiple Chinese provinces between December of 2019 and January of 2020. These isolates are likely ancestors of viral isolates recovered from patients in the USA, India, and Taiwan between January and March 2020. Interestingly, one isolate from Wuhan (LR757995.1) is part of the Group B cluster, supporting the hypothesis that multiple viral variants emerged in China before the start of the pandemic.

The Group C cluster was the only cluster that did not contain sequences from China. This cluster includes viral isolates collected from the USA, Italy, Australia, Sweden, Brazil, and South Korea between January and February of 2020. The absence of viral isolates from China and the increased genetic distance from Group A suggests that the emergence of these variants might have come from a second wave of transmission outside of China after the start of the pandemic.

### Evolutionary divergence in the genome of SARS-CoV-2

Once we reconstructed the phylodynamic of SARS-CoV-2 isolates obtained early during the pandemic event, we attempted to determine which nucleotide positions in the SARS-CoV-2 genome were related to the early divergence of this virus. Overall, the evolutionary rate of SARS-CoV-2 is 1.15 × 10^−3^ substitutions/site/year (95% HPD 7.41 × 10^−4^ -1.57 × 10^−3^), while pairwise analysis at nucleotide and amino acid levels revealed an average identity of 99.93-99.98% and 99.86-99.97%, respectively. Given the short divergence time, a high level of identity is to be expected; however, a few synonymous and non-synonymous substitutions were observed in the ORF1ab, S, OFR3a M, ORF8a, N and OFR10 genes (Figure 3A and 3B). When pairwise distances were calculated based on gene length, the highest levels of divergence were observed within genes ORF10 and ORF8a when considering synonymous and non-synonymous substitutions, respectively (Figure 3C and 3D).

**Figure 3.**
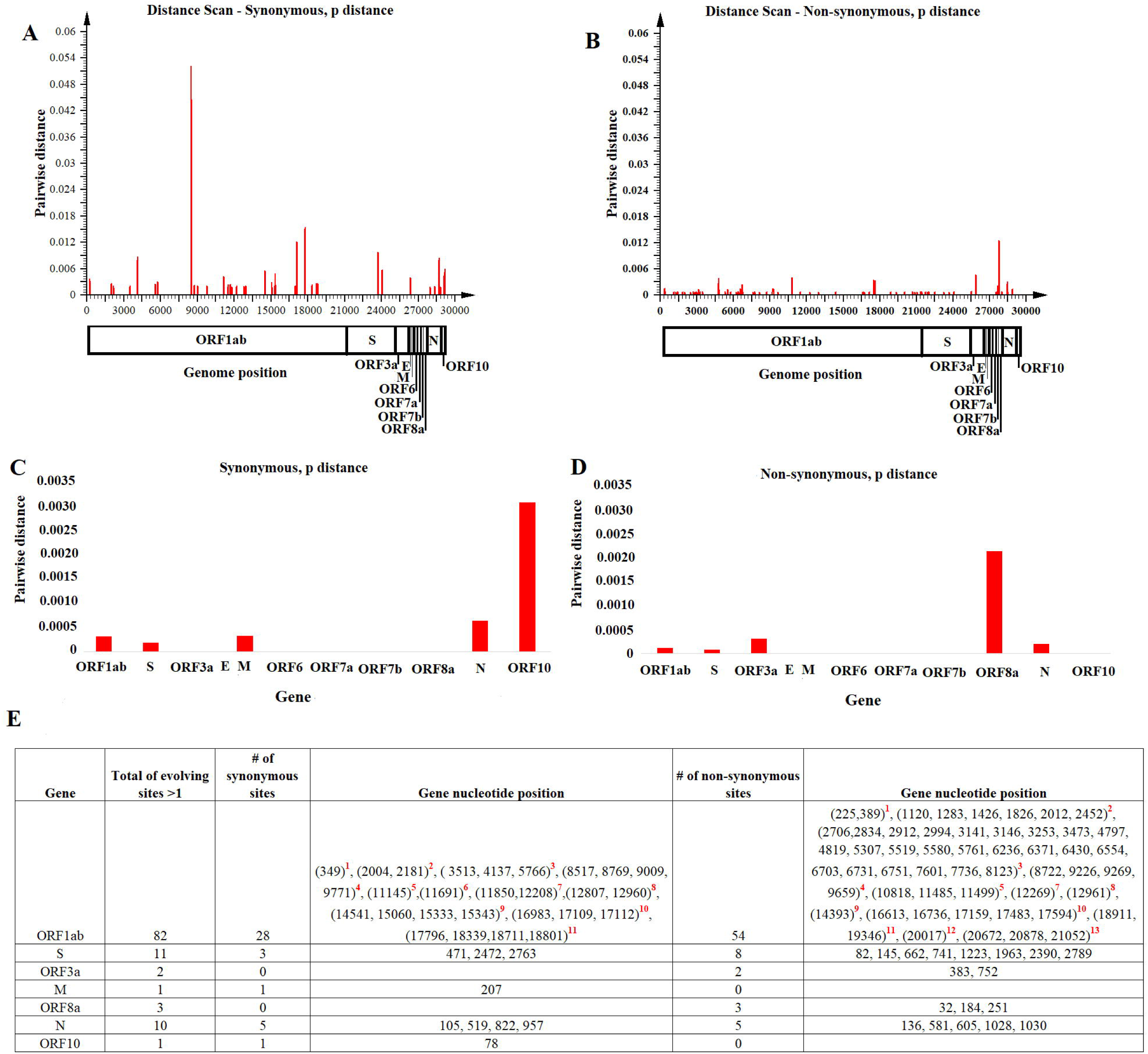
Pairwise distance analysis. Pairwise distance analysis at: A) synonymous and B) non-synonymous nucleotide sites was conducted using the program Sequence Distances (software SSE). Red bars represent pairwise distance comparisons using an sliding window of 50 nucleotides. Average nucleotide pairwise distance for different genes is shown at C) synonymous and D) non-synonymous sites. E) Fast-evolving synonymous and non-synonymous sites at each coding region are shown. For these sites evolutionary rates oscillated between 4.97 and 4.95. Red numbers represent nucleotides at : 1)Leader protein, 2) nsp2, 3) nsp3, 4) nsp4, 5) nsp6, 6) nsp7, 7)nsp8, 8)nsp10, 9) RNA independent polymerase, 10) helicase, 11) 3’to 5’ exonuclease, 12) endoRNAse, 13) 2’-O-ribose methyltransferase.

Also, the estimated per site evolutionary rate in the coding regions revealed that 98.85% of the sites in the genome are evolving at expected rates of evolution, while 1.15% of the sites are evolving faster than expected (Figure 3E). In this context, and consistent with the length of the OFR1ab gene, most of these synonymous and non-synonymous substitutions (82 sites) were distributed among different protein-encoding segments of this gene; the segment encoding nsp3 had the highest number of polymorphic sites (Figure 3E).

### Detection of purifying and diversifying selection

Once we identified fast-evolving positions within different genes of SARS-CoV-2, we used a combination of different algorithms centered on a codon-based phylogenetic framework to detect specific codons evolving under natural selection. Overall, no recombination events potentially affecting the results of these analyses were detected using the GARD algorithm.

Using SLAC we obtained a broad picture of the extent of natural selection acting upon the SARS-CoV-2 genome. We found an overall dN/dS ratio of 0.937 along the genome. In particular, 75 codons located within 5 genes (ORF1ab > S > N > ORF8a >ORF3a) showed evidence of increased fixation of non-synonymous mutations (dN/dS >1). Conversely, a small number of codons (35 codons) located within 5 genes (ORF1ab > N > S >M = ORF10) were accumulating a higher number of synonymous mutations (dN/dS <1). Interestingly, evaluation of dN/dS at the level of individual genes showed higher ratios for the ORF3a and ORF8 genes (Figure 4A).

**Figure 4.**
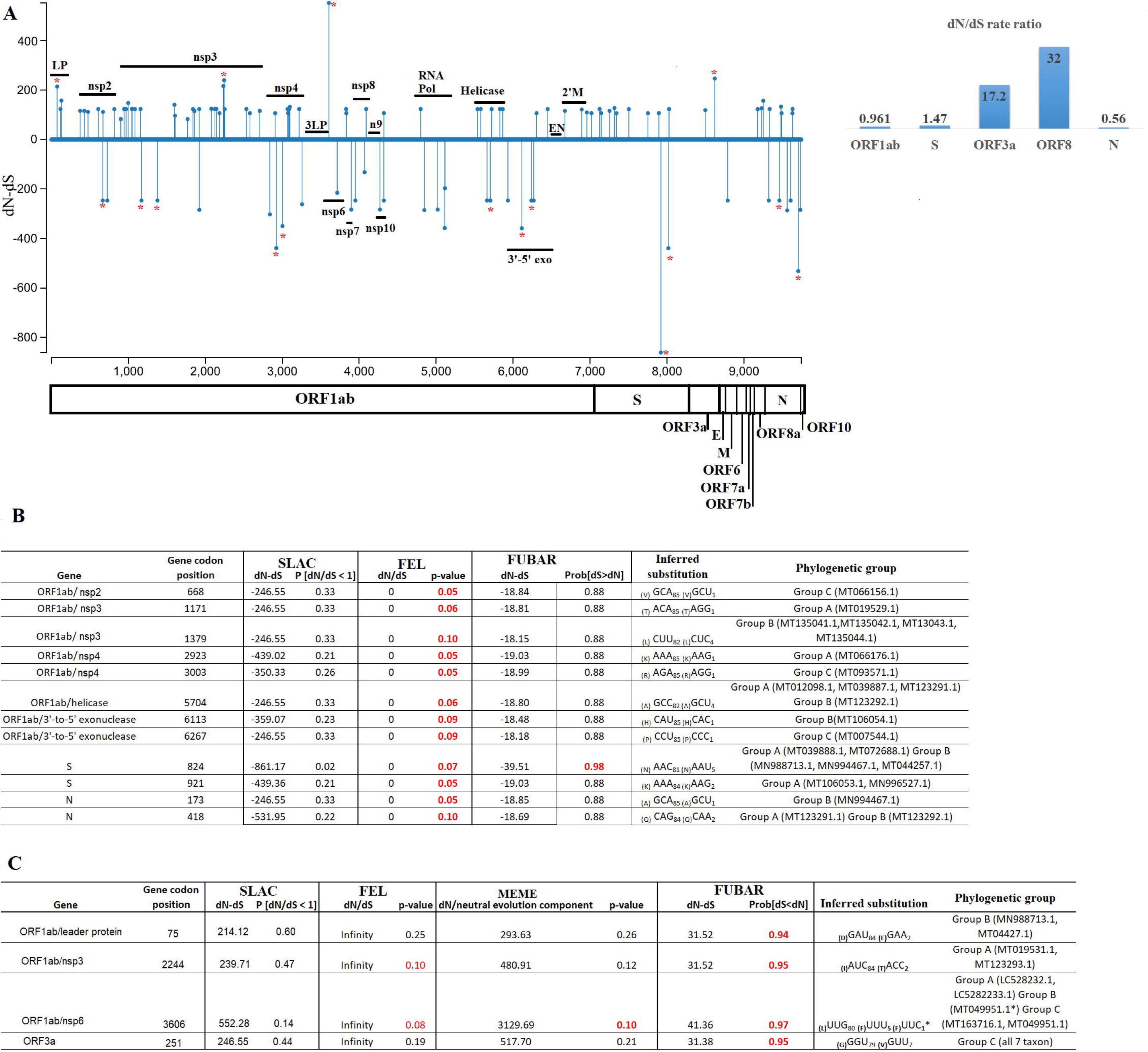
Diversifying and purifying selection on SARS-CoV-2. A) General overview obtained by SLAC analysis, showing the evolutionary rate (dN-dS or dN/dS) along the genome and at individual genes of SARS-CoV-2. Statistically significant codons were inferred by multiple evolutionary tests used in this study. Red asterisks represent codons with significant evidence for selection, Codons evolving at B) Purifying (negative) or C) Diversifying (positive) selection are shown numbers in red represent evolutionary tests with significant values according to the analysis :SLAC, FEL, MEME (p-value=0.1) and FUBAR (posterior probability =0.9). The criteria for considering a site positively or negatively selected was based on their identification by at least one of the tests. The Phylogenetic group column (assigned according with figure 1A) shows also the isolates carrying the substitutions. Abbreviations: LP (Leader protein), 3LP (3C-like proteinase), n9 (nsp9), 3’-5’ exo (3’-to 5’ exonuclease), EN (endoRNAse), and 2’M (2’-o-ribose methyltransferase).

Significant purifying (negative) selection was observed in 12 out of the 35 codons evolving at dN/dS <1 using the FEL (12 sites) and FUBAR (1 site) methods; the codons were located in the ORF1ab, S, and N genes (Figure 4B). At these codons, increased fixation of synonymous substitutions seems to be favoring the phenotypic preservation of SARS-CoV-2 at specific residues of the proteins encoded by these genes. By tracking these mutations within different isolates, we observed that these changes could explain the divergence of different viruses within different phylogenetic groups. In some cases, mutations were associated with multiple isolates, supporting the relevance of these findings.

On the other hand, evidence of diversifying positive selection on non-synonymous sites was detected in just 4 of the 75 codons evolving at dN/dS >1 in genes ORF1ab and ORF3a, with the FUBAR and FEL methods providing the highest power of detection (Figure 4C). Based on this analysis, these four sites appear to be evolving under pervasive diversifying selection.

Interestingly, the detection of diversifying selection at codon 3606 (nsp6) was significantly supported by three different tests. Also, the selection of this site was observed in isolates from all three phylogenetic groups, thus supporting the reliability of this finding. However, the conservative nature of the amino acid substitution at this site (L-F; BSM62= 0) suggests this may not affect the phenotype of SARS-CoV-2. The same pattern was observed in codon 75 (D-E; BSM62= 2).

Conversely, based on the nature of the substitution (G-V; BSM62= -3), diversifying selection at codon 251 of the ORF3a gene may produce a biologically relevant effect on the phenotype of SARS-CoV-2. The same situation was observed at codon 2244 (I-T; BSM62= -1). Interestingly, change at codon position 251 is highly conserved within isolates of group C, suggesting that this change might have promoted the divergence of this group.

### Detection of directional selection

To maximize the inference of potential sites experiencing positive selection, amino acid alignments of SARS-CoV-2 were analyzed using the DEPS algorithm. Overall, DEPS identified a total of 4 amino acid residues that are experiencing directional selection. Of these four residues, isoleucine (I) has the strongest bias, affecting 16 out of 19 sites evolving via directional selection (figure 5A).

**Figure 5.**
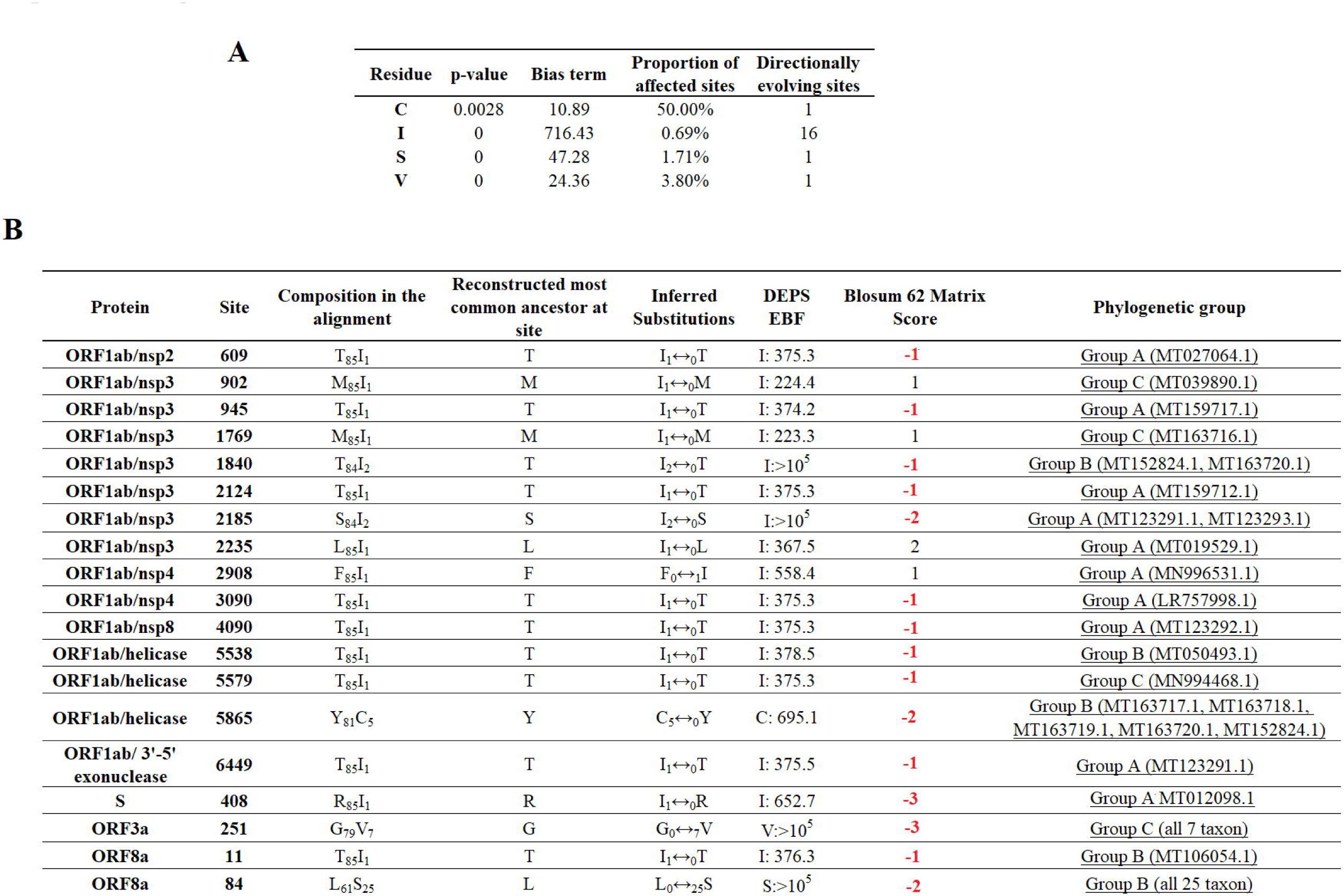
Directional selection analysis on SARS-CoV-2. A)An amino acid alignment was evaluated by DEPS and 4 different residues producing 19 directionally evolving sites in the proteome of SARS-CoV-2 are reported. P-values show the statistical significance of each residue considering a model test of selection versus not selection. Bias term: Alignment-wide relative rate of substitution towards target residue., Proportion of affected sites: Percentage of sites evolving under a directional model, versus a standard model with no directionality. Directionally evolving sites: Number of sites that show evidence of directional selection for focal residue. B) Description of 19 directionally evolving sites. Sites were detected by Empirical Bayesian Factor (EBF) considering a cut-off of 100 or more. Numbers in red represent replacements between amino acids with different properties. The Phylogenetic group column (assigned according with figure 1A) shows also the isolates carrying the substitutions.

The majority of selected sites were located in nonstructural proteins (nsp) encoded by the ORF1ab gene, with nsp3 accounting for the highest proportion (Figure 5B). Interestingly, a low proportion of the total number of predicted sites had a conservative amino acid substitution (residues at positions 902, 1769, 2235, and 2908), suggesting that the majority of substitutions may have an adaptive effect. In this context, it is remarkable that codon 84 of protein ORF8a is synapomorphic in all Group B sequences. Also, similar to previous algorithms, DEPS identified positive selection of residue 251 of ORF3a, supporting the potential significance of this site in the early evolution of SARS-CoV-2.

### Evidence of coevolution among sites

Finally, we attempted to find coevolutionary correlations between different codons within the genome that result in the positive selection of sites. Analysis by BMG produced evidence of 14 coevolving codon pairs; these interactions took place mostly within codons located within the ORFB1ab gene (Figure 6). Although most of the interactions were detected between nonsynonymous codons, coevolution between codons 4090-4269 and 818-4320 was detected by a synonymous substitution at one of the codons. Also, based on the nature of the amino acid replacement, just 6 of the 14 interactions produced a potential biological significance in the replacement. Interestingly, 8 of the 14 interactions appeared associated with sites evolving under some type of positive selection, suggesting that the selection of these sites might be the result of epistatic events (Figure 7).

**Figure 6.**
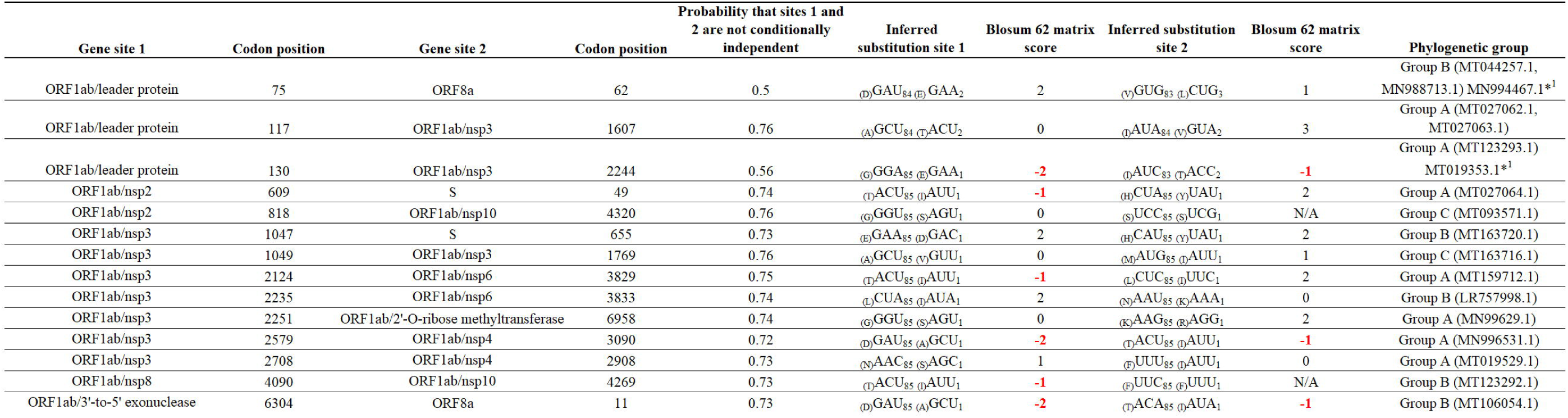
Coevolution between codon pairs in the genome of SARS-CoV-2. BMG analysis was conducted to detect coevolving codon pairs. Evidence of 14 coevolving codon pairs were detected and the specific locations of those in the genome of SARS-CoV-2 are presented. Posterior probability of pair associations was supported by Markov Chain Monte Carlo Analysis at cut-off of 50 or more. Numbers in red represent replacements between amino acids with different properties. The Phylogenetic group column (assigned according with figure 1A) shows also the isolates carrying the substitutions. *^1^ represents viral isolated where the changes were not detected.

**Figure 7.**
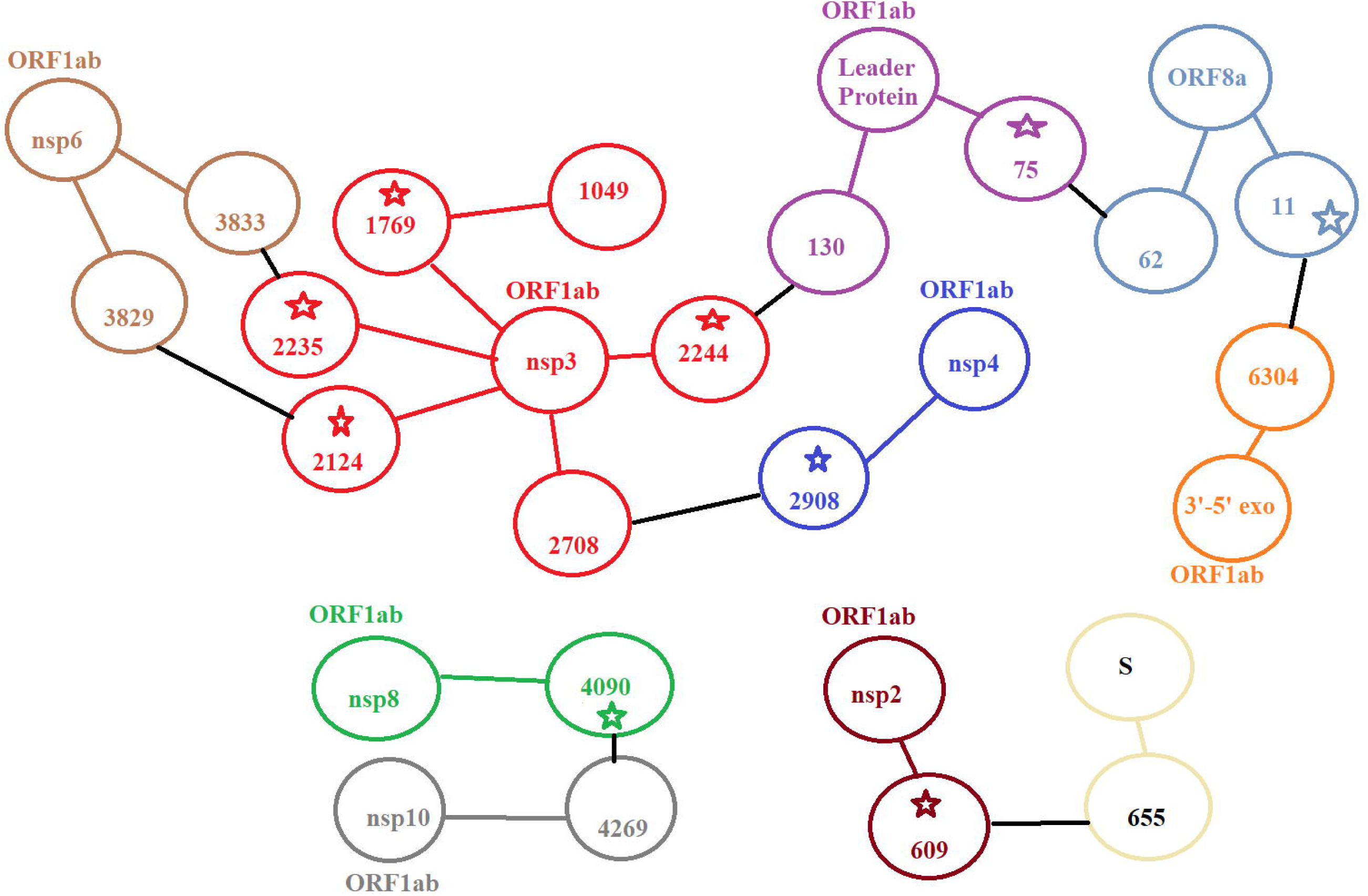
Potential epistatic network in the genome of SARS-CoV-2. Coevolving pair codons involving sites evolving under positive selection at different regions in the genome of SARS-CoV-2 are shown with stars, suggesting that the selection of these sites may be result of epistasis.

## Discussion

Herein, we evaluated the phylogenetic and evolutionary dynamics of SARS-CoV-2 during the first months of the pandemic event in 2020. Our phylogenetic analysis revealed the complex dynamic of the spread of infection throughout the world, suggesting that multiple viral variants might have emerged in China before the start of the pandemic event. The evolutionary rate calculated for SARS-CoV-2 in this study was consistent not only with previous reports for SARS-CoV (Salemi et al., 2004, Zhao et al., 2004), but also with the rate for other RNA viruses (Sanjuan et al., 2010), explaining the high levels of identity at nucleotide and amino acid levels calculated for SARS-CoV-2 in our study. In this context, the high conservation observed in the genome of SARS-CoV-2 early during the pandemic might also be attributed to the unique RNA correction machinery of coronaviruses (Ferron et al., 2018).

However, and despite the relative genome stability observed in SARS-CoV-2 at this stage of the pandemic, we were able to describe the existence of 3 phylogenetic groups. Interestingly, our evolutionary analysis supported the hypothesis regarding early divergence events produced during the pandemic. By using a combination of different evolutionary algorithms, we detected multiple codon sites that may be promoting the divergence of SARS-CoV-2. However, two primary considerations must be addressed regarding the biological relevance of multiple sites detected in this study. First, a considerable number of polymorphisms were detected in just one viral isolate, which might be a consequence of the small number of viral isolates available at the time we started this study. Another possibility is that some of the polymorphisms might be due to sequencing errors. Additionally, some positively selected sites resulted in conservative replacements in terms of the similar nature of these amino acids, suggesting that they may not have a potential biological effect on the phenotype of SARS-CoV-2. However, the last assumption must be taken with caution, since experimental work on Chikungunya virus has shown that a conservative replacement of aspartic acid (D) with glutamic acid (E) at residue 350 of the E1 protein is able to alter the affinity of a monoclonal antibody for this site (Tuekprakhon et al., 2018).

We are reporting three potentially relevant residues that may be driving the evolution of SARS-CoV-2 in human populations. Residue 251 of the ORF3a protein appears to be experiencing diversifying selection and might be related to the emergence of viruses in phylogenetic Group C. The early selection of this site might have a biological relevance since the ORF3a protein has been associated with virulence of human coronaviruses by controlling not only the expression of cytokines and chemokines but also inducing necrotic cell death (Shi et al., 2019).

Also, a residue located at position 84 of the ORF8a protein was found to be evolving under directional selection and might be related to the emergence of Group B. This protein has been implicated in viral pathogenesis by regulating the initial innate response (McBride and Fielding, 2012, Shi et al., 2019). Interestingly, ORF3a and ORF8a accessory proteins had the highest dN/dS, indicating that a positive selective pressure is being exerted on these proteins during the pandemic.

Based on the known roles in tropism and virulence associated with the helicase of MERS-CoV, we consider it important to mention codon 5865 (ORF1ab) that was found to be evolving under directional selection and might be related to the divergence of 5 isolates from Washington, USA, forming a sub-cluster in Group B.

Finally, our analysis of coevolution revealed some potential epistatic interactions that might be driving the evolution of SARS-CoV-2. This mechanism has been proposed to explain the emergence of an Ebola virus variant in 2014 (Ibeh et al., 2016), and its relevance in the evolution of coronaviruses should be explored in future studies. Also, it is interesting to mention that most of the co-evolving sites were located in nsp3; given the role of this protein in the virulence of coronaviruses (Fehr et al., 2015), this observation may be key in understanding the evolution of SARS-CoV-2. Furthermore, since two of the interactions detected by BGM were associated with synonymous mutations, the relevance of this type of substitution to viral fitness should not be underestimated, since selection of synonymous substitutions has been reported in other RNA viruses like VSV (Novella et al., 2004, Velazquez-Salinas et al., 2018).

Collectively, our results describe the early evolutionary events of SARS-CoV-2 during the current pandemic and the findings may support the hypothesis that different variants of SARS-CoV-2 with disparate levels of virulence might be circulating in the world. This possibility might have an important impact on public health and measures to control the pandemic. Subsequent studies using reverse genetics will be needed to confirm the relevance of our findings connecting specific residue substitutions with different virus phenotypes.

## Author Contributions

LV-S, and SZ conceived and designed the experiments. LV-S, SZ and SE performed the experiments. LV-S, SZ, SE, DG, IN, and MB analyzed the data. LV-S, SZ, SE, DG, IN, and MB wrote the manuscript.

## Funding

This work was performed under USDA Research Service CRIS Project No. 8064-32000-060-00D.

## Conflict of Interest Statement

The authors declare that the research was conducted in the absence of any commercial or financial relationships that could be construed as a potential conflict of interest.

## Acknowledgements

We would like to particularly like to thank Melanie Prarat for editing the manuscript. Also, we thank Dr. Peter Simmonds for the use of the SSE program used for pairwise distance analysis.

## Notes

### Competing Interest Statement

The authors have declared no competing interest.

## References

Andersen, K. G., Rambaut, A., Lipkin, W. I., Holmes, E. C. & Garry, R. F. 2020. The proximal origin of SARS-CoV-2. Nature Medicine.

CORONAVIRIDAE STUDY GROUP OF THE INTERNATIONAL COMMITTEE ON TAXONOMY OF, V. 2020. The species Severe acute respiratory syndrome-related coronavirus: classifying 2019-nCoV and naming it SARS-CoV-2. Nat Microbiol.

Delport, W., Poon, A. F., Frost, S. D. & Kosakovsky Pond, S. L. 2010. Datamonkey 2010: a suite of phylogenetic analysis tools for evolutionary biology. Bioinformatics, 26, 2455–7.

Drummond, A. J., Suchard, M. A., Xie, D. & Rambaut, A. 2012. Bayesian phylogenetics with BEAUti and the BEAST 1.7. Mol Biol Evol, 29, 1969–73.

Fehr, A. R., Athmer, J., Channappanavar, R., Phillips, J. M., Meyerholz, D. K. & Perlman, S. 2015. The nsp3 macrodomain promotes virulence in mice with coronavirus-induced encephalitis. J Virol, 89, 1523–36.

Ferron, F., Subissi, L., Silveira De Morais, A. T., Le, N. T. T., Sevajol, M., Gluais, L., Decroly, E., Vonrhein, C., Bricogne, G., Canard, B. & Imbert, I. 2018. Structural and molecular basis of mismatch correction and ribavirin excision from coronavirus RNA. Proc Natl Acad Sci U S A, 115, E162–E171.

Frost, S. D. W., Magalis, B. R. & Kosakovsky Pond, S. L. 2018. Neutral Theory and Rapidly Evolving Viral Pathogens. Mol Biol Evol, 35, 1348–1354.

Henikoff, S. & Henikoff, J. G. 1992. Amino acid substitution matrices from protein blocks. Proc Natl Acad Sci U S A, 89, 10915–9.

Ibeh, N., Nshogozabahizi, J. C. & Aris-Brosou, S. 2016. Both Epistasis and Diversifying Selection Drive the Structural Evolution of the Ebola Virus Glycoprotein Mucin-Like Domain. J Virol, 90, 5475–5484.

Kosakovsky Pond, S. L. & Frost, S. D. 2005. Not so different after all: a comparison of methods for detecting amino acid sites under selection. Mol Biol Evol, 22, 1208–22.

Kosakovsky Pond, S. L., Poon, A. F., Leigh Brown, A. J. & Frost, S. D. 2008. A maximum likelihood method for detecting directional evolution in protein sequences and its application to influenza A virus. Mol Biol Evol, 25, 1809–24.

Kosakovsky Pond, S. L., Posada, D., Gravenor, M. B., Woelk, C. H. & Frost, S. D. 2006. GARD: a genetic algorithm for recombination detection. Bioinformatics, 22, 3096–8.

Kumar, S., Stecher, G. & Tamura, K. 2016. MEGA7: Molecular Evolutionary Genetics Analysis Version 7.0 for Bigger Datasets. Mol Biol Evol, 33, 1870–4.

Mcbride, R. & Fielding, B. C. 2012. The role of severe acute respiratory syndrome (SARS)-coronavirus accessory proteins in virus pathogenesis. Viruses, 4, 2902–23.

Murrell, B., Moola, S., Mabona, A., Weighill, T., Sheward, D., Kosakovsky Pond, S. L. & Scheffler, K. 2013. FUBAR: a fast, unconstrained bayesian approximation for inferring selection. Mol Biol Evol, 30, 1196–205.

Murrell, B., Wertheim, J. O., Moola, S., Weighill, T., Scheffler, K. & Kosakovsky Pond, S. L. 2012. Detecting individual sites subject to episodic diversifying selection. PLoS Genet, 8, e1002764.

Novella, I. S., Zarate, S., Metzgar, D. & Ebendick-Corpus, B. E. 2004. Positive selection of synonymous mutations in vesicular stomatitis virus. J Mol Biol, 342, 1415–21.

Phan, M. V. T., Ngo Tri, T., Hong Anh, P., Baker, S., Kellam, P. & Cotten, M. 2018. Identification and characterization of Coronaviridae genomes from Vietnamese bats and rats based on conserved protein domains. Virus Evol, 4, vey035.

Poon, A. F., Lewis, F. I., Pond, S. L. & Frost, S. D. 2007. An evolutionary-network model reveals stratified interactions in the V3 loop of the HIV-1 envelope. PLoS Comput Biol, 3, e231.

Rambaut, A., Drummond, A. J., Xie, D., Baele, G. & Suchard, M. A. 2018. Posterior Summarization in Bayesian Phylogenetics Using Tracer 1.7. Syst Biol, 67, 901–904.

Salemi, M., Fitch, W. M., Ciccozzi, M., Ruiz-Alvarez, M. J., Rezza, G. & Lewis, M. J. 2004. Severe acute respiratory syndrome coronavirus sequence characteristics and evolutionary rate estimate from maximum likelihood analysis. J Virol, 78, 1602–3.

Sanjuan, R., Nebot, M. R., Chirico, N., Mansky, L. M. & Belshaw, R. 2010. Viral mutation rates. J Virol, 84, 9733–48.

Shi, C. S., Nabar, N. R., Huang, N. N. & Kehrl, J. H. 2019. SARS-Coronavirus Open Reading Frame-8b triggers intracellular stress pathways and activates NLRP3 inflammasomes. Cell Death Discov, 5, 101.

Simmonds, P. 2012. SSE: a nucleotide and amino acid sequence analysis platform. BMC Res Notes, 5, 50.

Sun, P., Qie, S., Liu, Z., Ren, J., Li, K. & Xi, J. 2020. Clinical characteristics of hospitalized patients with SARS-CoV-2 infection: A single arm meta-analysis. J Med Virol.

Tuekprakhon, A., Nakayama, E. E., Bartholomeeusen, K., Puiprom, O., Sasaki, T., Huits, R., Luplertlop, N., Kosoltanapiwat, N., Maneekan, P., Arien, K. K., Shioda, T. & Leaungwutiwong, P. 2018. Variation at position 350 in the Chikungunya virus 6K-E1 protein determines the sensitivity of detection in a rapid E1-antigen test. Sci Rep, 8, 1094.

Velazquez-Salinas, L., Pauszek, S. J., Stenfeldt, C., O’Hearn, E. S., Pacheco, J. M., Borca, M. V., Verdugo-Rodriguez, A., Arzt, J. & Rodriguez, L. L. 2018. Increased Virulence of an Epidemic Strain of Vesicular Stomatitis Virus Is Associated With Interference of the Innate Response in Pigs. Front Microbiol, 9, 1891.

Wan, Y., Shang, J., Graham, R., Baric, R. S. & Li, F. 2020. Receptor Recognition by the Novel Coronavirus from Wuhan: an Analysis Based on Decade-Long Structural Studies of SARS Coronavirus. J Virol, 94.

Weaver, S., Shank, S. D., Spielman, S. J., Li, M., Muse, S. V. & Kosakovsky Pond, S. L. 2018. Datamonkey 2.0: A Modern Web Application for Characterizing Selective and Other Evolutionary Processes. Mol Biol Evol, 35, 773–777.

Weiss, S. R. & Navas-Martin, S. 2005. Coronavirus pathogenesis and the emerging pathogen severe acute respiratory syndrome coronavirus. Microbiol Mol Biol Rev, 69, 635–64.

Wu, F., Zhao, S., Yu, B., Chen, Y. M., Wang, W., Song, Z. G., Hu, Y., Tao, Z. W., Tian, J. H., Pei, Y. Y., Yuan, M. L., Zhang, Y. L., Dai, F. H., Liu, Y., Wang, Q. M., Zheng, J. J., Xu, L., Holmes, E. C. & Zhang, Y. Z. 2020. A new coronavirus associated with human respiratory disease in China. Nature, 579, 265–269.

Zhao, Z., Li, H., Wu, X., Zhong, Y., Zhang, K., Zhang, Y. P., Boerwinkle, E. & Fu, Y. X. 2004. Moderate mutation rate in the SARS coronavirus genome and its implications. BMC Evol Biol, 4, 21.

